# Lipid order degradation in autoimmune demyelination probed by polarization resolved coherent Raman microscopy

**DOI:** 10.1101/105965

**Authors:** Paulina Gasecka, Alexandre Jaouen, Fatma-Zohra Bioud, Hilton B. de Aguiar, Julien Duboisset, Patrick Ferrand, Hervé Rigneault, Naveen K. Balla, Franck Debarbieux, Sophie Brasselet

## Abstract

Myelin around axons is currently widely studied by structural analyses and large scale imaging techniques, with the goal to decipher its critical role in neuronal protection. While there is strong evidence that in myelin, lipid composition and lipid membrane morphology are affected during the progression of neurodegenerative diseases, there is no quantitative method yet to report its ultrastructure in tissues at both molecular and macroscopic levels, in conditions potentially compatible with *in vivo* observations. In this work, we study and quantify molecular order of lipids in myelin at sub-diffraction scales, using label-free polarization resolved Coherent Anti Stokes Raman (PR-CARS), which exploits CARS sensitivity to coupling between light polarization and oriented molecular vibrational bonds. Importantly, the method does not use any *a priori* parameters in the sample such as lipid type, orientational organization and composition. We show that lipid molecular order of myelin in the mouse spinal cord is significantly reduced throughout the progression of experimental autoimmune encephalomyelitis (EAE), a model for multiple sclerosis, even in myelin regions that appear morphologically unaffected. This technique permits to unravel molecular-scale perturbations of lipid layers at early stage of the demyelination progression, while the membrane architecture at the mesoscopic scale (here about 100 nm) seems much less affected. Such information cannot be brought by pure morphological observation and opens new prospectives towards molecular-scale understanding of neurodegenerative diseases.

## I. INTRODUCTION

The multilamellar myelin sheath wrapped around axons plays a crucial role for efficient transmission of nerve impulses, by acting as an electrical insulator^1^. One of the most important features defining the functionality of myelin is the extraordinary ability of its components to self-assemble into a highly ordered and stable structure^2^. This self-assembly is due to lipids (*≈* 70% by dry weight) and proteins (*≈* 30% by dry weight) which by interacting with each other form a tightly packed multilayered membrane with complex structure and lipid phases. In neurological disorders, this compact structure is highly perturbed^3,4^, leading to dysfunctions of the central nervous system (CNS). Numerous studies of the sequence of the myelin sheath degradation process in the CNS have been reported^5,6^, mostly concentrating on a mice model of multiple sclerosis (MS), the experimental autoimmune encephalomyelitis (EAE). The reported general patterns are the enlargement of individual myelin sheaths, due to an interlamellar splitting and loss of bilayer adhesion, swelling, and finally the formation of vesiculated myelin debris^7,8^.

Although the large malformations of the myelin ultrastructure in the advanced stages of demyelination have been reported, its finer structural variations at early stages remain poorly characterized, preventing a complete understanding of the molecular mechanism of myelin degradation in neurodegenerative diseases. So far, imaging techniques such as Transmission Electron Microscopy (TEM)^1^ and small angle X-ray diffraction (XRD)^9,10^ are used to access finer scales, however, they are highly invasive for the sample itself and lacking of biochemical specificity is also an issue. In addition, there are growing indications that the morphological alterations occur in conjunction with molecular modifications e.g. lipid composition (charged vs. uncharged^11,12^, saturated vs. unsaturated^11,13^) or lipid phases^14–17^ in the myelin sheath, which may contribute to the loss of lipid membrane inter-layer adhesion, budding and vesiculation^18^.

Being able to image those fine structural or molecular alterations in the myelin sheath during degradation is thus of critical importance for a better understanding of the course of the demyelination progression. Among optical techniques that encompass Optical Coherent Tomography^19^, confocal reflectance^20^, nonlinear optics^21,22^, Coherent Anti-Stokes Raman Scattering (CARS) microscopy is particularly interesting for its label-free nature, its chemical specificity and its spatial resolution of a few hundreds of nanometers. In CARS, a pump field (frequency *ω*_*p*_) and a Stokes field (*ω*_*s*_) interact with the sample to generate an anti-Stokes field at frequency 2*ω*_*p*_ − *ω*_*s*_, which is enhanced when *ω*_*p*_ − *ω*_*s*_ is tuned to the resonant frequency of a Raman band^23–25^. By addressing lipid molecules by their CH vibration bonds, which are highly abundant in the myelin sheath, CARS microscopy has been used to image myelin in the mouse brain^26,27^, nerves of the peripheral nervous system^28,29^, as well as spinal cord tissues^13,14,27,30–34^ including during the progression of EAE^14,27,32^. Using CARS images, the commonly used parameter in the assessment of the myelin health by its morphology is the *g*-ratio, defined as the ratio between the inner and outer diameter of myelin fibers^14,28–30,33,35^. This parameter however does not give any information on the possible molecular alterations. Furthermore, association of CARS with confocal Raman microspectroscopy has evidenced a higher degree of lipid unsaturation at sites of degraded pathological myelin, a feature that is associated with higher lipid-packing disorder^13^. These findings suggest that lipid organization is affected in EAE tissues, even before any clear morphological sign of damage^13^.

In this prospect, polarization-resolved CARS (PR-CARS) has been developed, which exploits the sensitivity of CARS to the coupling between incident excitation polarizations and the orientation of CH bond vibrations in lipids^36^. In PR-CARS, the rotation of the incident pump and possibly Stokes linear polarizations generates a CARS signal modulation, which comes from the maximization of the CARS signal when the optical polarizations lie along the direction of the excited vibration modes^37^. In lipids PR-CARS imaging in myelin, where CH_2_ vibration modes are excited, this CARS modulation has been so far translated into a molecular order quantity supposing simple ”rigid lipid” models^14,29,30,34,38^. While being simplified, this model allowed to reveal changes of lipid organization due to demyelination in a murin model of Krabbe disease^38^, or in developmental myelination in the zebrafish spinal cord^34^. The ”rigid lipid” model however does not reveal a quantitative and physical picture of the orientational behavior of lipids in complex environments such as myelin. First, it supposes a unique lipid symmetry, ignoring possible variations in lipids conformations and composition. Second, it limits to the interpretation of angular order while another important information lies in the phase of the CARS signal modulation, which represents the average direction of molecular bonds within the optical focus of excitation. This orientation information brings insights into possible local deformations of the membrane at mesoscopic scales (~ 100 nm below the diffraction limit size), where membrane contour rupture or loss of lipid layer adhesion can occur for instance. Such scales are particularly important in the context of demyelination processes where lipid membranes are packed into multilayer structures. In this work, we use a generic approach by analyzing the complete information accessible from PR-CARS experiments, revealing both lipid order microscopic and mesoscopic scale behaviors in demyelination processes occurring in neurodegeneration. For this we use a more general symmetry decomposition supposing unknown orientational distribution of CH bonds^36,39,40^. By using such approach, we quantify the molecular order of CH bonds in myelin sheaths during active demyelination processes, as well as their averaged orientations, without any *a priori* knowledge on the lipid type, conformation or membrane shape. We show that PR-CARS signal modulation can inform on molecular scale disorder ultrastructure within the focal spot of optical resolution *~* 200 nm, but also at mesoscopic scales, which address modifications of molecular order over a distance of the image pixels size (typically *~* 100 nm). Performing PR-CARS analysis on spinal cord tissues taken from EAE mice at different stages of the disease, we show that early alterations of the myelin sheath occur at the molecular level, while the membrane architecture at the mesoscopic scale is seen to be particularly robust and mostly preserved until the formation of myelin debris. PR-CARS associated with such generic analysis is shown to be a potential tool for label free early detection of demyelinating disease progression.

## II. MATERIALS AND METHODS

### A. EAE induction and sample preparation

All experimental procedures were performed in accordance with the French legislation and in compliance with the European Community Council Directive of November 24, 1986 (86/609/EEC) for the care and use of laboratory animals. The research on animals was authorized by the Direction Départementale des Services Vétérinaires des Bouches-du-Rhône (license D-13 *−* 055 *−* 21) and approved by the National Committee for Ethic in Animal Experimentation (Section N14; project 87 *−* 04122012). C57B*l*6 mice were housed in cages with food and water *ad libitum* in a 12 h light/dark cycle at 22 *±* 1C.

EAE was induced in seventeen C57B*l*6 mice using MOG (35 *−* 55), a myelin oligodendrocyte peptide (75 *µ*g in CFA, a complete Freund’s adjuvant, containing 800 *µ*g of killed mycobacterium tuberculosis) by s.q. injection into 3 different locations (bilaterally over the femur, and at the base of the tail), and pertussis toxin (400 ng) by i.p. injection on day 0 and 2. Additionally, a total of six control C57B*l*6 animals were used. Three of them received injections of an antigen-free emulsion (CFA containing 800 *µ*g of killed mycobacterium tuberculosis), and pertussis toxin on day 0 and 2. First clinical signs typically arised 13 *−* 14 days post-induction and were translated into clinical scores as follows: 0, no detectable signs of EAE; 0.5, tail weakness; 1, complete tail paralysis; 2, partial hind limb paralysis; 2.5, unilateral complete hind limb paralysis; 3, complete bilateral hind limb paralysis; 3.5, complete hind limb paralysis and partial forelimb paralysis; 4, total paralysis of forelimbs and hind limbs (mice with a score above 4.5 to be killed); and 5, death.

For the *ex vivo* sample preparation, groups of animals were sacrificed when reaching predetermined clinical scores for subsequent imaging of the myelin content in the dorsal spinal cord. Spinal cords were dissected, post-fixed overnight at 4C and rinsed 3 times in phosphate buffer saline (PBS) prior to embedding in a 3% agarose matrix to ease handling and 3D positioning under the microscope. Care was taken to align the dorsal surface of the spinal cord with the upper surface of the agarose cube. Note that the fixation method did not affect the results presented in this work. A few measurements in fresh tissues were performed that revealed very similar data.

### B. PR-CARS imaging

PR-CARS imaging (represented in Fig. 1a) was done using an OPO femtosecond laser source (80MHz 150fs, Coherent, Santa Clara, CA) with excitation wavelengths *λ*_*p*_ = 830 nm and *λ*_*S*_ = 1087 nm, temporally synchronized and spatially overlapped on the sample plane. The corresponding excited resonances correspond to lipid vibrations around 2850 cm^−1^ with a bandwidth 150 cm^−1^, dominated by the CH_2_ and CH_3_ stretching modes. The total power delivered to the sample is in the range of 10 mW, focused by a water immersion objective (40X/1.15W Nikon, Tokyo, Japan), which provides an optical resolution of about 250 nm. Imaging is performed in the epi inverted microscope geometry, using the dichroic mirror T770SPXR (Chroma Tech. Corp., Bellows Falls, VT). Scanning mirrors rate is 20 *µ*s per pixel, over 30 *µ*m size images (200 *×* 200 pixels). The nonlinear signal collected by the objective is filtered using a shortpass filter (ET750sp-2p8, Chroma Tech. Corp., Bellows Falls, VT) before being detected by a PMT (R9110, Hamamatsu, Shizuoka, Japan) in the spectral range of 650 nm using a bandpass filter 655AF50 (Omega Optical, INC.). Polarization imaging is done by rotating the linear polarization angle of the incoming beam in 10 steps over the range 0-170(Fig. 1b), using a NIR achromatic half wave plate (AHWP05M-980, Thorlabs, Newton, NJ) mounted in a motorized rotation mount (PR50CC, Newport, Irvine, CA). An image is recorded at each polarization step as explained in ^41^. This results in a polarimetric image stack (Fig. 1b) where each pixel contains a PR-CARS modulated response.

**FIG. 1.**
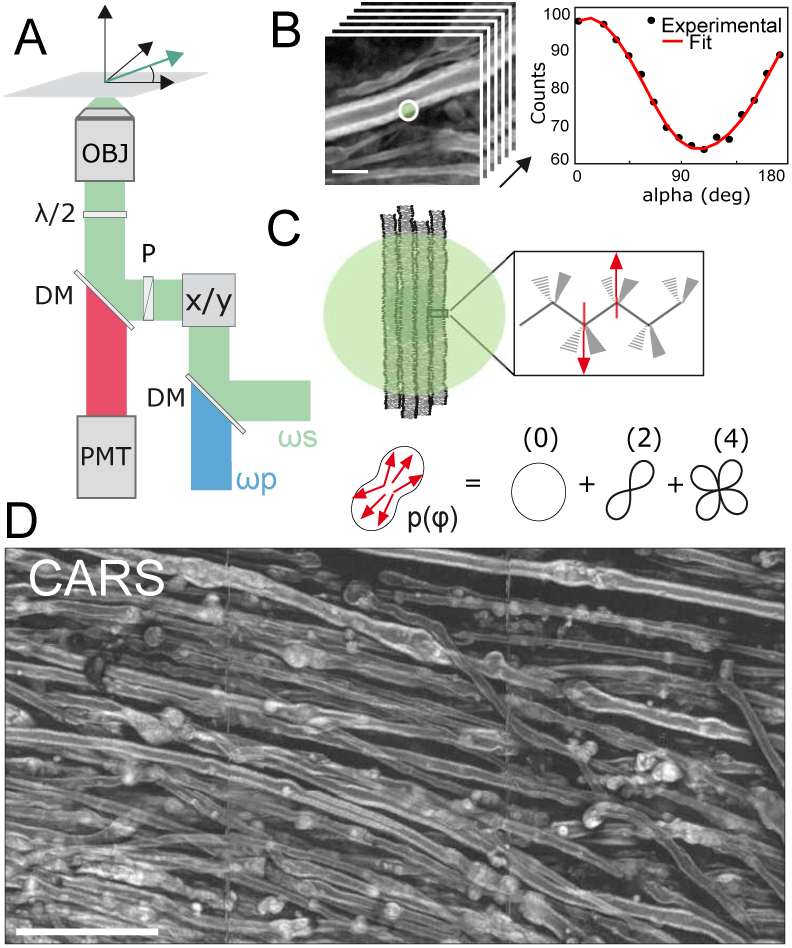
(*a*) Schematic view of the PR-CARS microscope. The vector *E* represents the direction of linear polarization of *ω*_*s*_ and *ω*_*p*_(*α* angle), rotated by a half-wave plate (*λ*/2). DM, dichroic mirror; x/y, galvanometric mirrors; P, polarizer; PMT, photomultiplier tube; OBJ : objective. (*b*) Example of PR-CARS data (left panel) recorded in one pixel (right panel) of the myelin sheath region (control sample), averaged over the diffraction limit size (schematically represented by a circle). Scale bar: 5 *µ*m. (*c*) Top: Schematic of the multilamellar lipid membrane in the focal spot (represented as a circle) and zoom on a lipid showing the dominant nonlinear bond dipoles (red arrows) probed by PR-CARS. Bottom: distribution function *p*(*φ*) read-out from PR-CARS (see text), as a sum of orders of symmetry. (*d*) Large scale CARS image (total intensity summed over 18 input polarization angles) of a spinal cord tissue (score 1.5). Scale bar: 50 *µ*m.

### C. Data processing and analysis

The quantification of molecular order from PR-CARS measurements was previously described in^39,40^. Briefly, both linearly polarized incident pump (*E*_*p*_) and Stokes fields (*E*_*s*_) are rotated with a variable angle *α* relative to the sample plane horizontal axis *X* (Fig. 1a,b). The PR-CARS response is then analyzed to retrieve orientational parameters from the lipid assembly present in the focal volume of the objective. The PR-CARS signal is derived from a bond additive model where individual CH molecular bonds (represented as arrows in Fig. 1c) form the excited vibrational modes. These molecular bonds are organized into a 2D effective distribution function *p*(*φ*), that can be decomposed over its symmetry orders (Fig. 1c):

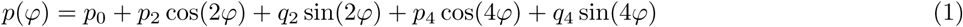

where *p*_0_ is the isotropic contribution to the angular distribution, (*p*_2_, *q*_2_) are its second order symmetry components and (*p*_4_, *q*_4_) its fourth order symmetry components. This decomposition allows to avoid any hypothesis on the measured vibrational modes population, which could come from different types of lipids and different lipid conformation.

The molecular bonds add-up coherently to form the CARS nonlinear coherent radiation, which intensity is decomposed on a circular basis (See Supporting Material):

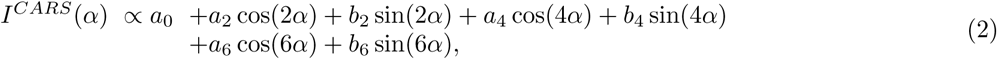

The (*a*_*n*_, *b*_*n*_) coeffcients are directly calculated from the recorded polarization-dependent image stack, by projection on circular basis functions.

In the PR-CARS data analysis, the (*p*_*n*_, *q*_*n*_) parameters are then deduced from the (*a*_*n*_, *b*_*n*_) coeffcients following the relations derived in^39,40^ (See Supporting Material). To define the properties of the distribution function parameters measured for each pixel of the PR-CARS image, we introduce the magnitudes (*S*_2_, *S*_4_) and orientations (*φ*_2_, *φ*_4_) of the second and fourth symmetry order contributions respectively:

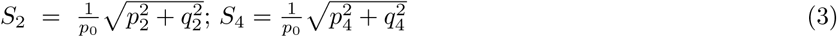

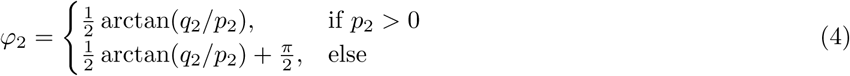

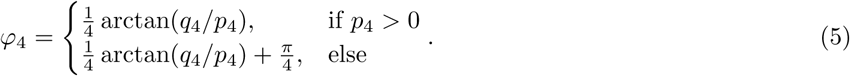

The anisotropic nature of an angular distribution function is mainly quantified by its contribution *S*_2_ and main orientation *φ*_2_, while *S*_4_ and *φ*_4_ quantify the shape (smooth versus sharp) of this distribution. It is convenient to rewrite *S*_4_ as a symmetric component that is of cylindrical symmetry along *φ*_2_ (denoted 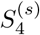) and an asymmetric contribution (denoted 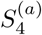)^42^:

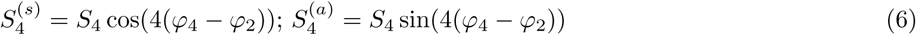

In particular if the distribution is of cylindrical symmetry, then *φ*_2_ = *φ*_4_ or *φ*_2_ = *φ*_4_ *±* 45°, and 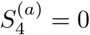. Theoretical analyzes on modeled angular distribution functions show that the parameters 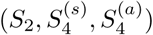 permit to discriminate between different shapes of molecular angular distributions^39,42^.

In practice, the selection of the relevant pixels to be analyzed is performed on the basis of the total CARS intensity image obtained by summing for each pixel the signal obtained for all angles *α*. To perform intensity thresholding based on criteria discussed below, the PR-CARS intensity is averaged over 3 × 3 pixels and only pixels which total intensity is above 75% of the threshold are considered. Pixels corresponding to the signal from the myelin sheath and its different morphological variations are manually selected using a polygon selection tool of Matlab (The MathWorks, Natick, MA). For the estimation of the *g*-ratio parameter in straight myelin sheaths, the ginput function of Matlab (The MathWorks, Natick, MA) is used. We made sure that the myelin sheaths regions selected to calculate the molecular order parameters corresponds to *g*-ratio values within a range of 0.05 relative to the mean *g*-ratio of the given population. The retrieved molecular order parameters 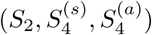 are indicated with their mean and standard deviation values, measured over a large collection of myelin sheaths regions for each given population. Statistical significance was determined using ANOVA tests performed over those populations with (*\** : *p* ≤ 0.05) considered as statistically significant; (** : *p* ≤ 0.01) as highly statistically significant; (*** : *p* ≤ 0.001) as extremely statistically significant.

## III. RESULTS

Fig. 1d shows a typical CARS image of myelin sheaths in a mouse spinal cord, imaged at about 20 *−* 30 *µ*m below the spinal cord surface with linear polarization excitation along the spinal cord axis. The CARS signal is dominated by symmetric and antisymmetric CH_2_ stretching vibrations of hydrocarbon chains from lipids (2845 cm^−1^ and 2885 cm^−1^ respectively) which are the major components of myelin. CH_3_ stretching vibration of proteins (2930 cm^−1^) also contribute to a lesser extent, since they constitute 15 *−* 30% of the weight of the myelin sheath^13,14^. Fig. 1d and the zoomed view of Fig. 1b show longitudinal sections of lipid multilayers of about 3 *−* 5 *µ*m in total diameter, wrapped around axons. The myelin morphology is seen to be strongly affected when the EAE score disease increases, from straight myelin tubular structures to swelled features (appearing at score 1) coming from vesiculated myelin, and finally to the formation of debris with spherical shapes of various sizes, coming from disrupted myelin detached from the axon (with a major proportion at scores 2.5 and 4).

### A. Molecular order of lipids in the myelin sheath

A PR-CARS polarization analysis has been performed on zoomed-in regions of the myelin sheath structure, using a high sampling pixel size of 100 nm. Fig. 2 shows typical PR-CARS results in a spinal cord of a control tissue.

**FIG. 2.**
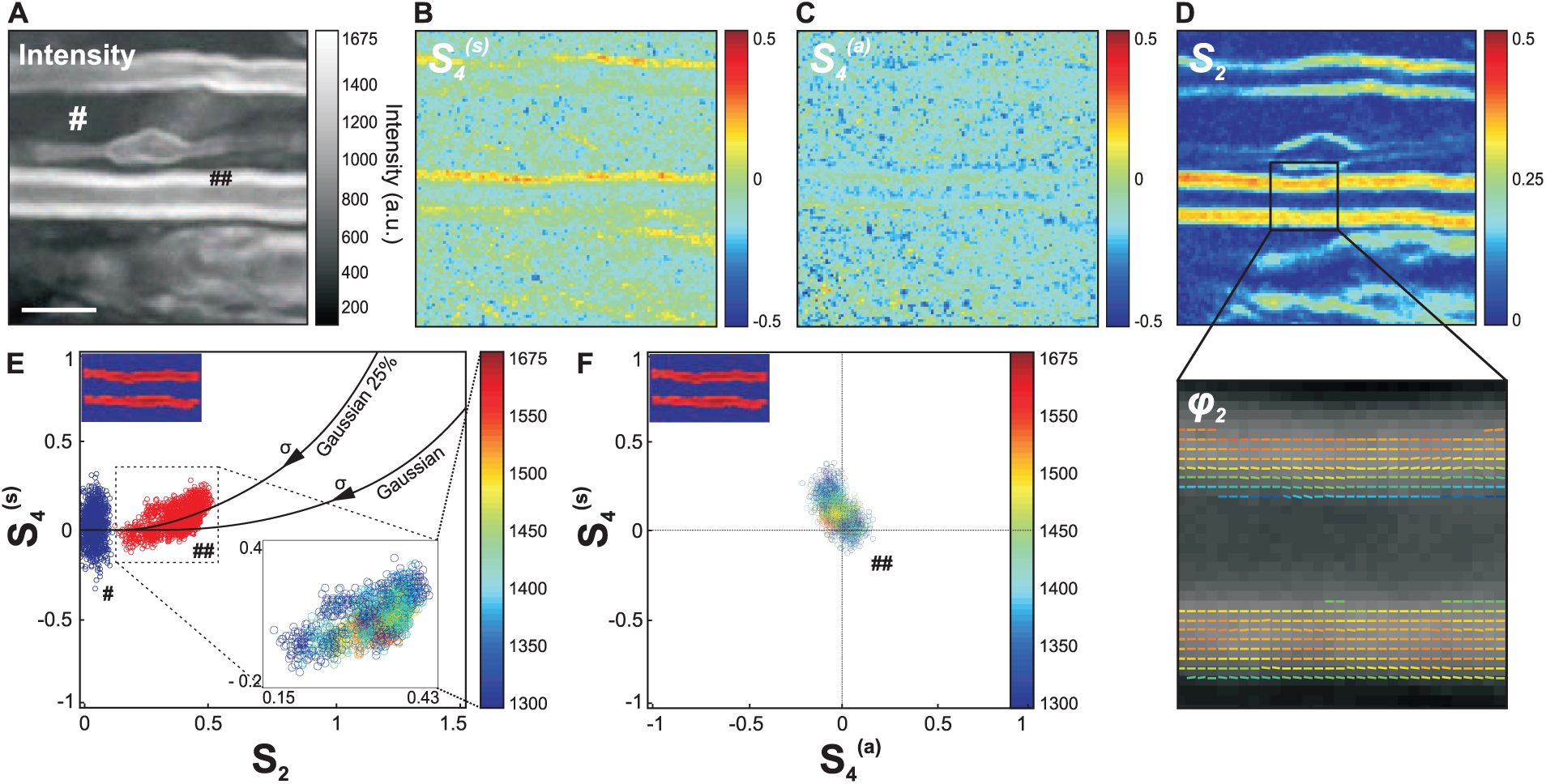
(*a*) PR-CARS (centered at 2845 cm^−1^) total intensity image (sum over 18 input polarization angles) of the myelin sheath from a spinal tissue (control sample). Scale bar: 5 *µ*m. (*b*) Corresponding 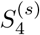 map. (*c*) 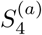 map. (*d*) *S*_2_ map. For those maps (not for data analysis) the PR-CARS intensity is averaged over 3 *×* 3 pixels. A region of the *S*_2_ map is zoomed-in to make more visible the complete orientational information, represented by a stick oriented with the *φ*_2_ angle with respect to the horizontal axis, and which color is *S*_2_. (*e*) Experimental values of *S*_2_, 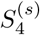 taken at intensities above threshold 1300 analog signal value in the myelin sheath (*♯♯* in the intensity image (a)) and at lower intensity signals from a surrounding background (*♯* in the intensity image (a)). For the myelin sheath region, the inset image is color coded in intensity. Solid lines: theoretical 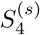 values for a Gaussian distribution of increasing width *σ*, and Gaussian superimposed with an isotropic distribution of proportions 25 %. (*f*) 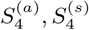 scatter plot following similar color coded intensity scale as in (*e*).

The interpretation of the measured 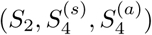 values depicted in Fig. 2b-d requires a rigorous control of measurement conditions, in order to avoid bias and error sources (see Supporting Material). Briefly, the intensity level is set above a threshold that guaranties standard deviations of *S*_2_ below 0.02, and of 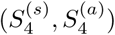 below 0.08 (see Fig.S1 in the Supporting Material). A residual but not significant intensity-dependent bias is also systematically removed (see Fig.S1 in the Supporting Material). The *S*_2_ values additionally suffer from a bias due to the presence of non-resonant background^40^, which can be tolerated in relative comparison studies and therefore not removed. Another source of bias can also occur due to the out-of plane orientation of the measured distributions, *S*_2_ values being at their maximum when the distribution lies in the sample plane. This effect is excluded by a control of the plane of focus at the equatorial plane of the myelin sheath (see Fig. S2 in the Supporting Material). At last, birefringence is seen to lead to negligible bias of the deduced values at the depths of measurement used in this analysis (see Fig. S3 in the Supporting Material). Note that scattering from the spinal cord tissue can be also neglected, previous studies have indeed shown that effects due to the detected multiply scattered light in epi geometry do not affect the polarization CARS signals as long as the total intensity (non-analyzed detection) is recorded^43^.

The *S*_2_ image is strongly contrasted (Fig. 2d), which is consistent with the tight packing of lipids in the multilamellar structure of myelin, while the background is isotropic (*S*_2_ = 0). The *φ*_2_ image (Fig. 2d) shows additionally that the general orientation of CH bond dipoles in the equatorial plane is along the myelin tubular axis, as expected from lipids chains oriented radially in a concentric axonal structure. The fourth order symmetry of the CH bonds distribution provides additional information, since 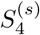 is clearly above the surrounding background (Fig. 2b,e). The 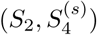 experimental values are plotted together as a scatter graph in Fig. 2e to provide a finer interpretation of the CH bonds orientational distribution in the lipid multilamellar structure. Importantly, the measured order parameters are seen to not depend on the intensity (Fig. 2e), which excludes experimental artifact related to noise level. Continuous lines on this graph represent theoretical dependencies expected from different orientational distributions of CH bonds, that span similar regions as experimental values. For a Gaussian distribution of width *σ*, *S*_2_ is expected to grow with decreasing *σ*, without considerable increase of 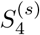 except at low widths (*σ* < 80°). The measured 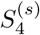 values are however higher than expected from a Gaussian distribution. This can be explained by the presence of an isotropic angular population superimposed to a Gaussian function^42^, which could occur from small disordered regions in the focal volume, such as sub-diffraction size vesicles. Fig. 2e depicts different cases of distributions composed of *η* % of an isotropic population superimposed with (100 − *η*) % of a Gaussian population, showing that the measured values correspond to *η* ≈ 25 % (accounting for the fact that *S*_2_ is probably slightly underestimated due to the presence of non-resonant background). At last, 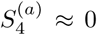 (Fig. 2c,f), confirming the cylindrical symmetry of the orientational distribution of bonds in the lipid organization. In what follows, *S*_4_ will be often used as a shortened notation for 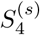, and 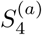 will be discarded.

Overall, the average values < *S*_2_ >= 0.33 *±* 0.05 and < *S*_4_ >= 0.10 *±* 0.07 obtained in the myelin sheath depicts a molecular order contained in an angular width of about *σ* ≈ 80°. Interestingly, this order is not far from that observed in model membranes; PR-CARS performed in multilamellar vesicles (MLVs) made of pure lipids indeed show that < *S*_2_ >≈ 0.22 in dioleoylphosphatidylcholine (DOPC) MLVs and < *S*_2_ >≈ 0.40 in dipalmitoylphosphatidylcholine (DPPC) MLVs, the latter exhibiting a higher liquid order phase transition temperature (see Fig.S4 in Supporting Material). The presence of several lipid types and proteins in the myelin sheath visibly does not enlarge considerably the CH bonds angular distributions as compared to MLVs, confirming the tight organization of lipids along the axons. Looking closer at standard deviations however, the orientational behavior of CH bonds exhibits a higher level of heterogeneity than in MLVs (see Fig.S4 in Supporting Material), which is coherent with their diversity and the complexity of the inter-lamellar interactions in myelin.

### B. Relation between lipid packing orientational order and myelin morphology

During demyelination, the compact multilamellar structure of myelin undergoes swelling and degradation, until the formation of detached debris. In this section we investigate if there is a correlation between the morphology of the myelin sheath and lipid order measured by PR-CARS, regardless of the score of the EAE disease progression. A more quantitative comparison between scores will be given in the next section.

PR-CARS measurements were performed at different stages of the EAE disease progression in mice, from control to score 4. CARS imaging was first performed to localize areas on fixed spinal tissues, chosen randomly over several mm distances. Second, PR-CARS was performed in several small ROIs (typically 20 × 20 *µ*m size) at various places. CARS images permitted to determine five categories which are considered as representative of the different characteristic morphological features observed during the demyelination process, depicted in Fig. 3a: ”normal” myelin has a compact structure, with visible straight borders forming a tubular structure around the axon; ”swollen” myelin defines more loose membrane formations; ”blebbed” denotes a later stage of swelling, which forms a more pronounced protrusion outside of the straight myelin sheath structure; ”debris” denotes a lipid multilamellar spherical structure from vesiculated myelin, that has detached from the myelin sheath. Such debris, reported in the literature^7,8^, are found in two categories: “non filled debris”, which show no lipid signal at their center, and filled debris that are pure lipids volumetric structures. We observed an increased amount of damaged or vesiculated myelin with the progression of the disease as previously reported^13,32^.

**FIG. 3.**
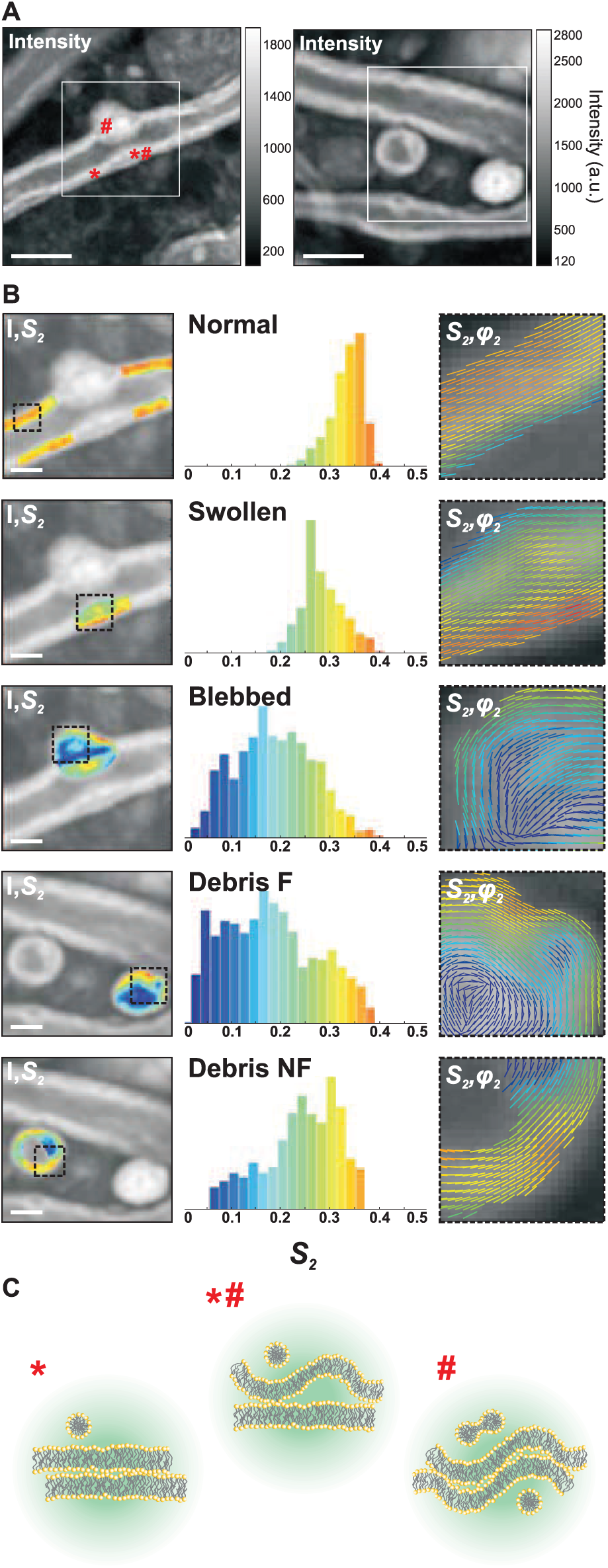
PR-CARS analysis of the different characteristic morphological features visible in the demyelination process. (*a*) Total intensity images taken from clinical score 1 (left) and 2 (right). Symbols (*\*, ♯, *♯*) refer to different membrane locations. White square region of interests (ROIs) are used for statistical analysis in (b). Scale bars: 5 *µ*m. (*b*) Left: *S*_2_ maps (superimposed on the intensity image) of characteristic features of myelin degradation in white ROIs of (a): ’healthy’ myelin, ’swollen’, ’blebbed’ and products of degradation: ’debris not filled (NF)’ and ’debris filled (F)’. Middle: corresponding histograms of *S*_2_ values, taken within the dashed square ROIs in (b). Right: combined images of *S*_2_ and *φ*_2_, showing *S*_2_ as a colorscale and *φ*_2_ as the orientation of sticks for each measured pixel. (*c*) Schematic morphological interpretation of the myelin multilayer (enlarged) in positions in (a) corresponding to (*) ’healthy’, (*\*♯*) ’swollen’ and (*♯*) ’blebbed’ myelin. The circular region represents the focal spot (not to scale).

For each category we measured molecular order of CH bonds, deducing (*S*_2_, *S*_4_) from PR-CARS data. We did not observe significant changes of *S*_4_ in this study, therefore this parameter is omitted in what follows. Fig. 3b (left) depicts zoomed-in images of the *S*_2_ parameter measured on each type of category. The *S*_2_ histograms (Fig. 3b, middle) collected for each of those characteristic regions illustrate the heterogeneity of molecular order in degraded myelin. At last, the mean orientation (*φ*_2_) are represented in these regions (Fig. 3b, right) depicts the spatial properties of such molecular order in healthy and damaged regions.

A few key observations can be made: (1) the average *S*_2_ values decrease from normal (*S*_2_ *≈* 0.30) to swollen (*S*_2_ ≈ 0.25) and blebbed (*S*_2_ ≈ 0.17) myelin, indicating loss of molecular scale order. Fig. 3c depicts such situations schematically, supposing that this disorder originates purely from morphological nanometric scale perturbations; (2) in blebbed and debris structures, *S*_2_ is more heterogeneous than in normal structures, with lower molecular order at the inner edge of the membrane; (3) molecular order in debris is preserved at their edge, with only inner membranes perturbed with high disorder. Lipid order in the periphery of not-filled debris is rather high as compared to filled debris, similarly as what has been observed in DPPC MLVs in which the filled structures exhibits a lower molecular order (see Fig.S4 in Supporting Material). In not-filled debris, CH molecular order can even reach values measured in normal myelin sheaths; (4) the overall orientation of CH bonds, illustrated by (*S*_2_,*φ*_2_) images, seems to follow the membrane macroscopic direction whatever the disorder, even for low *S*_2_ regions where the molecular scale disorder is high. This indicates that at the mesoscopic scale represented by the pixel size (here 100 nm), no strong modification of the membrane structure is found, while disorder is still rather occurring at a molecular/nanometric scale, averaged within the focal sport size. Overall these observations show that the transformation of normal, compact myelin structure into the final debris results in important local nanometric rearrangements of the multilamellar membrane structure, which seem to be able to persist over mesoscopic scale distances.

The values of molecular order measured here for a given region, stay valid over a large population of regions. Measurements performed on 359 regions (on average 70 regions per category) are summarized in global histograms in Fig. 4a. Independently of the stage of the EAE disease, normal myelin exhibits the highest degree of lipid packing (< *S*_2_ >= 0.32), which corresponds to the smallest aperture *σ* of the effective distribution function ≈ 80. The degradation of lipid packing in the myelin lamellar structure of swollen regions is manifested by a decrease down to < *S*_2_ >= 0.23, which correspond to an aperture *σ* of ≈ 100. In blebbed myelin, the averaged global order goes down to < *S*_2_ >= 0.14, which corresponds to an aperture *σ* of ≈ 120. Concerning myelin degeneration products, the outer membrane of not-filled debris exhibit an averaged order of < *S*_2_ >= 0.25, which is higher than for filled ones < *S*_2_ >= 0.17. All cases show significant differences between each others, except for not-filled debris which resemble normal myelin.

**FIG. 4.**
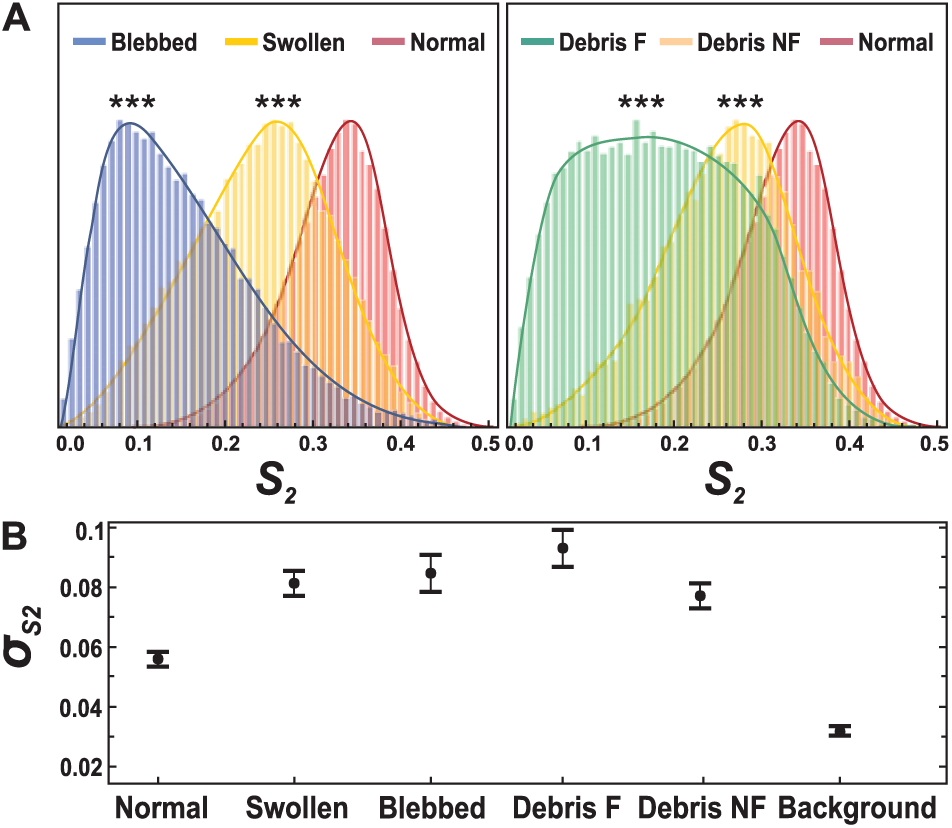
(*a*) General normalized histograms (over all measured pixels) of characteristic features of myelin degradation plotted for all measurements, regardless the clinical score of the EAE disease (about 1000 pixels per regions). Number of regions measured: normal myelin (N = 136), swollen (N = 91), blebbed (N = 82), debris filled (F) (N = 30), debris not-filled (NF) (N = 20). *\*\*\** : *p* ≤ 0.001. (*b*) Corresponding standard deviations *σ*_*S*_2__ within each region, taken per category. Error bars are standard deviation values over all regions measured.

The heterogeneity of molecular order within these regions is seen to increase for degraded myelin, as compared to normal myelin (Fig. 4b). The standard deviation of *S*_2_ (*σ*_*S*_2__) shows that in normal myelin, a low standard deviation is measured (*σ*_*S*_2__ = 0.06). This value is not noise-limited (as visible in the low background value where *σ*_*S*_2__ = 0.03), but still characteristic of very homogeneous lipid packing. From normal to debris, this standard deviation increases with *σ*_*S*_2__ = 0.08 in swollen, blebbed myelin and not-filled debris, and *σ*_*S*_2__ = 0.09 in filled debris. The filled debris regions indeed exhibit a very wide histogram, mostly due to the fact that in these structures the inner membrane shows very disordered characteristics.

### C. Evolution of lipid organization in the myelin sheath with the progression of EAE

The different myelin morphological features described above, which characterize demyelination, were visible in all samples independently of the EAE clinical score, even though their occurrences varied from score to score. In this Section, we evaluate how *S*_2_ in these different regions, depends on the clinical scores of EAE. This should allow an assessment of the finer evolution of myelin microscopic-scale properties during neurodegeneration. For each measured score, two spinal cords have been imaged with equal number of regions per spinal cord, except for score 4 where only one spinal cord was measured. Additionally, injection of complete Freund’s adjuvant (CFA sample) was produced in order to evaluate the consequence of the induction of an inflammation which is not specific to EAE.

First, normal myelin can be classified depending on the thickness of its sheath. For this we quantified the *g*-ratio of all measured normal-looking myelin sheaths, defined by the ratio between the inner to the outer diameter of the myelin membrane (Fig. 5a). Values of *g*-ratio are found to be in the range from 0.3 to 0.65, which is consistent with what has been reported in the literature^14,30,33^. When plotting the *g*-ratio values versus < *S*_2_ > values measured in all studied cases independently of the EAE score, only a slight correlation between the two parameters is visible (Fig. 5b). We also plotted *g*-ratio values for the different clinical scores of EAE (Fig. 5c). The variation of *g*-ratio values was seen to be very large even inside a given score population (as visible by the large standard deviations). Note that the high value measured for CFA samples might be due to myelin swelling, as a result of the immune response to the injection. Contrary to previous works^28,44^ that observed the thinning of myelin sheath (decrease of the *g*-ratio), we do not observe drastic trend of dependence between *g*-ratio and normal-appearing myelin condition during the progression of EAE.

**FIG. 5.**
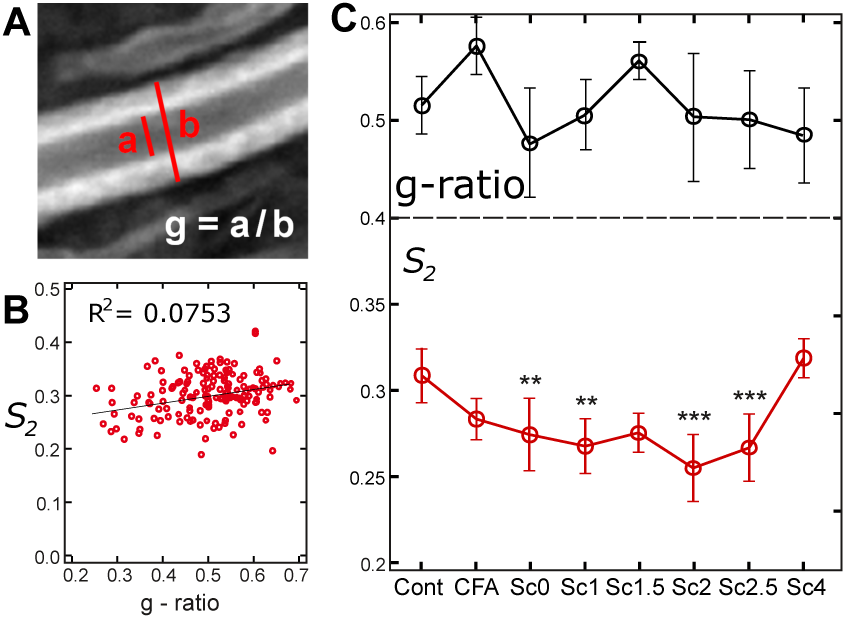
(*a*) Measurement of the *g*-ratio in the equatorial plane of a myelin sheath, using the total intensity image (*a*: inner diameter, *b*: outer diameter, *g*-ratio is the ratio between the inner and outer diameters). (*b*) Mean < *S*_2_ > values of all measurements versus *g*-ratio, with linear interpolation and correlation coefficient of the fit. (*c*) Top: mean *g*-ratio values versus the EAE score. Bottom: *S*_2_ values versus different scores of EAE, for the myelin regions of normal morphology (see text). Statistical analysis is shown relative to the control sample: * : *p* ≤ 0.05, ** : *p* ≤ 0.01, *** : *p* ≤ 0.001. Number of regions considered in the analysis: control (N = 26), CFA (N = 30), score 0 (N = 28), score 1 (N = 17), score 1.5 (N = 9), score 2 (N = 22), score 2.5 (N = 19), score 4 (N = 11). Error bars are standard deviation values.

In contrast, more pronounced variations were observed in *S*_2_ values measured in myelin sheaths of normal morphology for different EAE clinical scores (Fig. 5c). *S*_2_ in normal myelin morphologies decreases indeed progressively from 0.33 *±* 0.02 (control sample) down to 0.27 *±* 0.02 (score 2.5). All values show statistically significant changes as compared to the control and CFA sample, except scores 1.5 and 4 which are poorer in statistics. Note that score 4 contains a very low number of healthy-looking myelin sheaths, therefore the high *S*_2_ values measured are probably originating from the few robust myelin sheaths that survived the progression of the disease. These data show overall that molecular order in the myelin sheath of normal morphology decreases with the clinical signs of the EAE disease. Note that this trend is also visible when taking all myelin morphology types together, however in this case the *S*_2_ population might contain regions dominated by isotropic disorder present in myelin blebs and debris, making the error margins larger. Interestingly, the CFA sample shows no significant difference with the control sample, which suggests that pure immune reaction due to the injection does not provoke modifications of lipid order in myelin. In contrast, the observed loss of molecular order in EAE cases is thus seen to be specific to the disease.

## IV. DISCUSSION

Measuring molecular order in the myelin sheath by PR-CARS clearly permits to assess features in tissues that are not visible by pure (non PR) CARS imaging. The measurements performed highlight in particular that in the EAE demyelination process, the lipid multilayers organization of the myelin sheath is perturbed at a molecular/nanometric scale (as seen on *S*_2_ images), especially in the inner membrane region of the myelin sheath and from normal to swelled/blebbed myelin. This brings new elements of information in the understanding of lipid membrane behaviors in neurodegenerative diseases, which has been a topic of strong interest for decades. It has been reported that in the EAE pathology, different factors can contribute to the demyelination process, that cover molecular composition modifications and nanometric scale morphological changes. Even though PR-CARS microscopy cannot distinguish between those two factors, quantitative elements can be gained from the measured modifications of lipid molecular order. First, lipid composition changes have been reported in EAE models but not intensively studied. The existing observations have shown that lipids integrity (charge, degree of saturation) is altered to lead to an increase in membrane fluidity, and to a decrease of myelin adhesion that provokes vesiculation^11,13^. Our PR-CARS analyzes on the other hand have shown that going from a liquid disorder phase (typically DOPC MLV at room temperature) to liquid order phases (typically DPPC at room temperature) can shift the measured order parameter *S*_2_ from 0.2 to 0.4. Observations in the myelin sheath show modifications along the same range of magnitude, which could partly be explained by local modifications of the lipid composition. Second, the membrane morphology of myelin is likely to be affected at the nanometric scale, as known from X-ray diffraction^45^ and electron microscopy studies^1^. Numerical models in PR optical microscopy have shown that increasing a single lipid membrane curvature can lead to significant changes in the read-out of the *S*_2_ parameter. Typically, a 10 nm height change within the diffraction limited spot of 200 nm leads to an increase of *S*_2_ of 0.1^46^. Reported local folding in demyelination processes, using X-ray diffraction, show that the highest efficiency of the myelin sheath to transfer an electrical impulse is reached when the lipid bilayers are highly compact against one to another, with a spacing of ≈ 3−4 nm between lipid layers^45^. An increase of spacing between bilayers (≈ 6 − 7 nm) has been seen to occur during the swelling process^45^. Our data suggest a decrease of *S*_2_ from normal to swollen of about 0.1, that could be consistent with such a small morphological modification. It is in particular likely that the PR-CARS measurements probes perturbations from well parallel stacked layers in normal myelin (* in Fig. 3a) to partly folded inter-layer membrane regions in swollen myelin (*\*♯* in Fig. 3a), and finally to completely folded multi-layer membrane regions in blebbed myelin (*♯* in Fig. 3a). Note that the fact that lipid orders in not-filled versus filled debris behave similarly as in DPPC MLVs tends to favor interpretations in the direction of modifications of the nanometric-scale membrane morphological changes. It is indeed probable that membrane folds are decreased due to higher membrane tensions, as already observed in cell membranes^46^.

More information can been gained from the averaged orientation of the CH bonds organization (*φ*_2_ images). Interestingly, even though molecular order can be strongly affected locally, the averaged orientation of molecular bonds still follows the macroscopic membrane direction. This means that even though the perturbation occurs at the molecular scale (lipids composition and interaction changes) or at the nanometric scale (sub-diffraction membrane folds), the membrane cohesion is preserved at the mesoscopic scale ( 100 nm pixel size) (Fig. 3b). Long persistence lengths of *φ*_2_ values are clearly visible whatever the degree of nanometric scale disorder in the local membrane, which could also explain the capacity of such local disruptions to create longer scale damage events (swelling, blebbing), that extend spatially at up to micrometers distances and ultimately lead to debris.

A striking feature of this study is that in apparently normal myelin sheaths, even though the macroscopic morphology is similar from score to score, molecular scale lipid disorder increases with the EAE disease progression. This result emphasizes the potential of PR-CARS to reveal features that are not visible at a macroscopic scale, which is particularly interesting in the context of the monitoring of the evolution of a pathology. Note that normal myelin can be easily identified regardless of the clinical score, and could thus serve ultimately as a convenient and robust assessment of the disease. This study also shows that in the context of EAE, the more traditionally used *g*-ratio factor, which is based on macroscopic-scale morphological information, seems less robust. Previous works have reported the use of *g*-ratio as an assessment of a neuronal condition following crush lesion^28,44^ or chemically induced demyelination^29,47^. In the first case, the thinning of myelin sheath (decrease of the *g*-ratio) was observed as a result of demyelination for the nerves of straight morphology, such as seen in the peripheral nervous system. In spinal cord or brain samples, however, axon morphologies can depart from straight features, especially at pathological states where a number of shape abnormalities are detected^27^. In such cases the measurement of *g*-ratio can be more delicate and easily biased. In the second case, the assessment of myelin health by the *g*-ratio upon a chemically induced demyelination seems to be very different from that seen in autoimmune induction (EAE). In chemically induced demyelination, the myelin sheath enlarges entirely and *g*-ratio increases as a result of degradation of internal membranes, while in EAE the degradation of myelin due to internal swelling and splitting of membranes seem to appear more locally, leaving some parts of the myelin sheath intact.

## V. CONCLUSION

In this work we have shown the possibility to probe, by PR-CARS, lipid orientational organization at the molecular level within the myelin sheath of mice spinal cord tissues, in the frame of demyelination and neurodegeneration. Modifications of lipid order have been found in myelin structures that seem morphologically unaffected, emphasizing the possibility of the technique to decipher processes that occur at early stages of demyelination. The higher degradation of lipids order have been found in the processes of swelling and blebbing, especially at the inner interface of the myelin sheath, an effect that is strong enough to induce long range deformations of the whole myelin sheath. This supports early findings which addressed local lipid modifications by biochemical analyzes, and advantageously complements such studies by imaging, which provides a way to address those questions potentially *in vivo*. Although the data are at this stage not spectrally resolved within the CH-based lipids vibration bands, for instance to discriminate saturated versus unsaturated lipids, future directions can be envisioned to study more precisely the origin of the observed phenomena by adding chemical specificity, at the expense however of lower signal levels. Ultimately if applied *in vivo*, this methodology could serve as a early stage detection for demyelinating diseases progression.

## VI. AUTHOR CONTRIBUTIONS

P.G. performed the experiments, prepared all MLV samples and analyzed data; A.J. and F.D. prepared all spinal cord samples; F.-Z. B. developed data analysis tools; N.B., H.B.A, J.D., P.F., and H.R. contributed to analysis tools and experimental developments; S.B. designed research and analytic tools; P.G., S.B. and F.D. wrote the manuscript.

## VII. ACKNOWLEDGMENTS

We thank D. Côté and A. Gasecka (Université Laval, Quebec, Canada) for providing the spinal cord slices used for birefringent measurements described in the Supporting Material. This work has been supported by CNRS, contracts ANR-15-CE19-0018-01 (MyDeepCARS), ANR-10-INBS-04-01 (France-BioImaging), ANR-11-INSB-0006 (France Life Imaging), and Conseil Regional Provence Alpes Côte d’Azur. F.-Z.B. received a scholarship from the Erasmus Mundus Doctorate Program Europhotonics (grant no. 159224-1-2009-1-FR-ERA MUNDUS-EMJD).

## VIII. SUPPORTING MATERIAL

### A. PR-CARS data analysis

The calculation of the CARS polarized (PR-CARS) response from the lipid assembly present in the focal volume of an objective relies on a bond additive model where individual CH molecular bonds add-up coherently to form a nonlinear susceptibility tensor *γ* for each vibration mode, responsible for the CARS nonlinear coherent radiation. These bond dipoles directions e with orientations (*θ*, *φ*) in the macroscopic frame form a macroscopic nonlinear susceptibility tensor 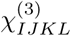 of the lipid assembly in the macroscopic 3D frame (*X, Y, Z*). It has been shown that since the incident optical field is polarized in the (*X, Y*) plane, the read-out of the CARS signal occurs from a reduced 2D macroscopic tensor^39^:

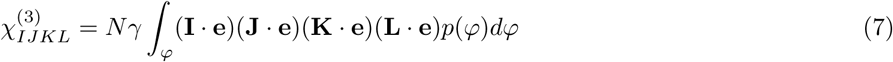

where (*I, J, K, L*) = (*X, Y*). *N* is the molecular bonds density, is the molecular vibration tensor norm, (I *·* e) refers to the projection of the e vector on the macroscopic frame axis I, and *p*(*φ*) is the orientational distribution function of e vectors averaged on their out-of plane orientation (*θ*)^39^. The nonlinear response finally depends on the function *p*(*φ*), a 2D effective distribution function that can be decomposed over its symmetry orders read by the PR-CARS process:

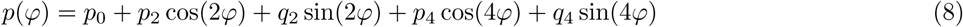

where *p*_0_ is the isotropic contribution to the angular distribution, (*p*_2_, *q*_2_) are its second order symmetry components and (*p*_4_, *q*_4_) its fourth order symmetry components. Relations between the macroscopic nonlinear susceptibility terms 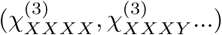 and the angular distribution function parameters ((*p*_2_, *q*_2_), …) can be found in^39^.

In the PR-CARS analysis, both incident pump field (*E*_*p*_) and Stokes field (*E*_*s*_) are linearly polarized and rotated with a variable angle *α* relative to the sample plane horizontal axis *X*. The CARS intensity can be written as^39^:

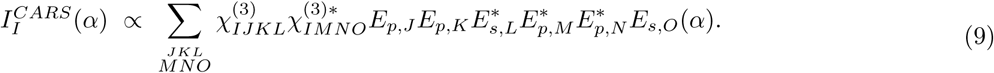

where *E*_*p,J*_ (*α*) is the projection of the pump incident field on the axis J in the sample plane. In the PR-CARS data analysis, the total CARS intensity is decomposed on a circular basis:

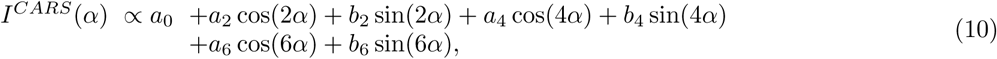

where the (*a*_*n*_, *b*_*n*_) coefficients are directly calculated from the recorded polarization dependent image stack by projection on circular basis functions.

The (*p*_*n*_, *q*_*n*_) parameters are deduced from these coefficients following the relations^39,40^:

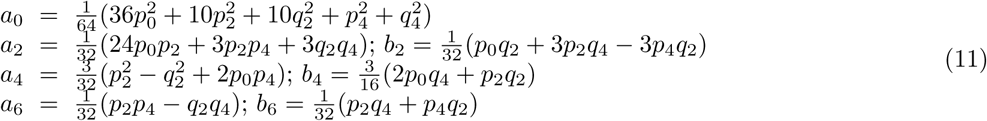

### B. Sources of error and bias in PR-CARS measurements

Stringent control of the PR-CARS signal conditions are required in order to ensure the validity of the measured order parameters. In this part the different sources of error and bias are investigated.

#### 1. Noise

First, *S*_2_, *S*_4_ values are only collected for high-enough intensity values, that guaranty sufficiently low bias and low noise (Fig. S1). Above 1300 analog signal value, standard deviations are of 0.02 for *S*_2_ and 0.08 for *S*_4_. This intensity value will be set as a threshold for all quantitative studies. Above this threshold, we also observe that the *S*_2_ bias is negligible. Concerning *S*_4_, the residual bias (which does not surpass 0.15 with the chosen intensities) is systematically removed from all measured values, accounting for the local pixel intensities and the reference noise-intensity dependence depicted (Fig. S1). In conclusion by using intensity thresholding, the measured parameters are relevant for statistical analysis and hence for comparison between different myelin regions and scores.

**FIG. S1.**
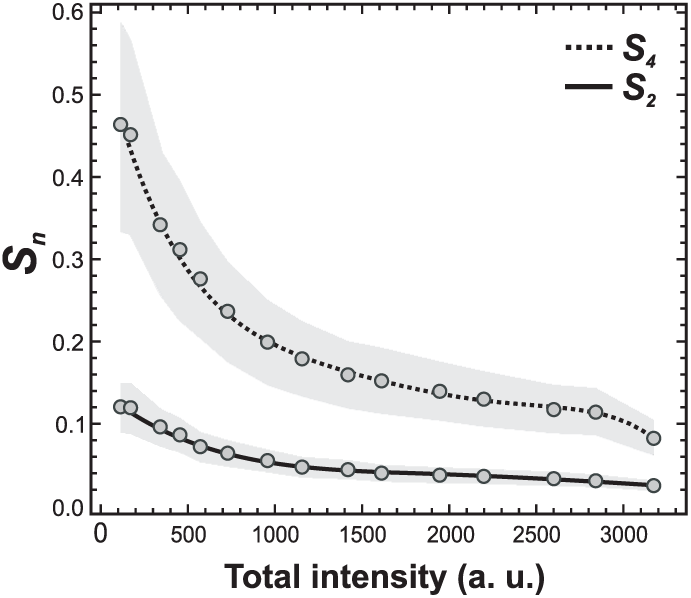
*S*_2_ and *S*_4_ dependence on the CARS intensity signal (summed over 18 input polarization angles) in the isotropic medium. The shaded region represents the standard deviation of the mean *S*_2_ and *S*_4_ values.

#### 2. Non-resonant background

Second, some care has to be taken regarding the existence of non-resonant background in the data. Considering our previous studies performed on model multilamellar lipid membrane vesicles, it is obvious that the non-resonant Four Wave Mixing (FWM) signal can lead to an underestimation of *S*_2_^40^. The difference between measured values and background-free ones (measured in absence of non-resonant background by Stimulated Raman Scattering (SRS)) have been estimated to be about 0.2 in multilamellar vesicles (MLVs) made of dipalmitoylphosphatidylcholine (DPPC), *S*_4_ being almost un-affected^40^. In this work we display raw data, however it is worth noting that the background-free *S*_2_ values have to be increased by an offset, which is most probably not higher than 0.2, considering that MLVs data are very close to values measured in myelin (see below).

#### 3. Out of plane mean orientation

Third, (*S*_2_, *S*_4_) values depend on possible out-of-plane orientations of the observed distributions. All the models developed in this work assume that the molecular angular distributions are lying in the sample plane. In order to properly report structural properties of myelinated axons, one needs to ensure that the PR-CARS measurement is performed at the equatorial plane of the axon/myelin tube. Any out-of-plane orientation of the observed distribution can indeed bias the results interpretation by adding a component which tends towards isotropy. In order to estimate how the (*S*_2_, *S*_4_) order parameters depend on this effect in the myelin sheath, we performed PR-CARS measurements at several focusing planes (*Z* steps every 0.5 *µ*m) crossing the whole 3D tubular structure of the myelin sheath from bottom to top (Fig. S2). *S*_2_ seems to be robust with negligible modification over a defocus of ±5 *µ*m around the equatorial plane. This means that as long as the myelin sheath looks visually in focus (which is the case for this defocus range), the *S*_2_ measurement is reliable. Moreover its standard deviation does not depend on Z. *S*_4_ on the other hand is very robust to a defocus change, which is consistent with the fact that defocus changes the apparent disorder of the distribution, not its shape. The only noticeable changes of *S*_4_ are a slight increase at the largest depth of focus, however in this region the intensity also drops down, which might increase the bias on *S*_4_.

**FIG. S2.**
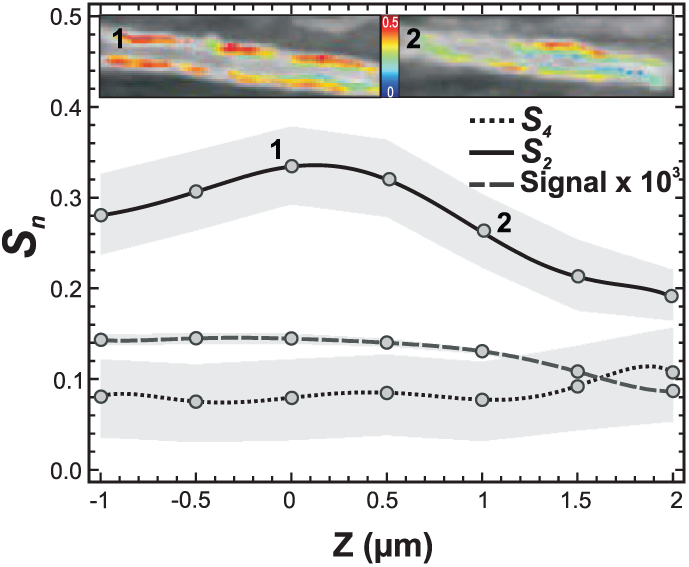
*S*_2_ and *S*_4_ mean values measured on a myelin sheath as a function of the sample depth Z. The shaded regions indicate the standard deviation of the order parameters (over all pixels in selected regions along the myelin sheath, according to an intensity thresholding of 75% of the maximum intensity measured in the image). Z= 0 corresponds to the depth of the myelin tube equatorial plane. Insets: *S*_2_ images after intensity thresholding, at Z= 0 (point 1) and 1 *µ*m above (point 2). Both images are superimposed with the CARS intensity images in grayscale.

#### 4. Birefringence

At last, the measured *S*_2_ values can also be underestimated if the sample exhibits birefringence. Birefringence acts indeed as a distortion for excitation fields’ polarization state, making them elliptic and tilted relative to their expected directions^48^. Birefringence is known to be non-negligible in the myelin sheath, due to the tight and regular arrangement of lipids^49^. We estimated the birefringence phase shift in myelin sample slices of thicknesses of about 30 *µ*m in order to extrapolate the expected birefringence in the measured spinal cord tissues (Fig. S3). The slices were cut after extraction and fixation in the sagittal plane using a vibratome, and placed in phosphate-buffered saline (PBS).

The measurement of birefringence follows a method presented in^37,39,50^ (Fig. S3). A laser beam (*λ* = 800 nm) is focused on the sample surface using the same microscope as used in PR-CARS. The incident linear polarization is rotated with an angle *α* (relative to the horizontal sample plane direction *X*) from 0 to 170 with a step of 10. The transmitted intensity modulation is recorded on a photomultiplier, placed after a polarizer of horizontal orientation (e.g. along the *X* direction of the sample plane, denoted (*X, Y*)). As a control in a non-birefringent sample, we performed the same experiment next to the spinal cord slice tissue in PBS. The resulting intensity polarization response along the *X* and *Y* analyzer directions can be written as quantities proportional to:

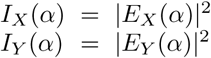

where *E*_*X,Y*_ are the transmitted field components along the *X* and *Y* directions respectively. Here only *I*_*X*_ (*α*) is measured. If birefringence occurs locally in a sample of given thickness, along a given optical axis direction Θ (typically the local direction of lipid chains in a thick multi-layer membrane), then the fields’ components along this axis and its perpendicular direction will be phase-shifted by a value Φ. Before reaching the birefringent medium, the optical electric fields are written (*E*_*X*_, *E*_*Y*_) = (cos *α*, sin *α*) in the laboratory frame. The birefringence imposes a phase shift Φ between the component of the field along the slow optical axis of the birefringent medium (denoted *X*_*b*_) and its fast axis (denoted *Y*_*b*_), such as:

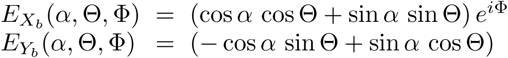

**FIG. S3.**
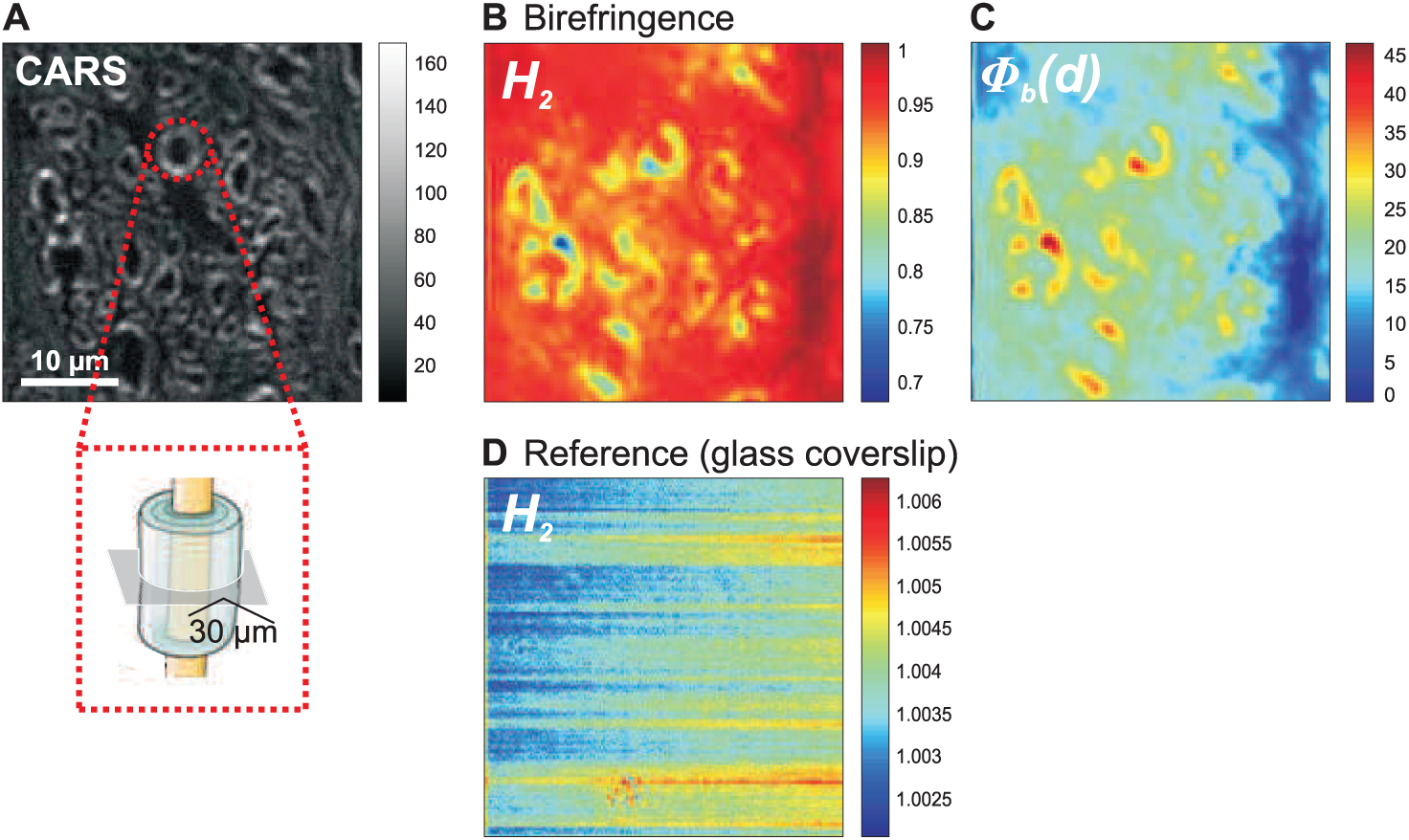
(*a*) CARS image (2845 cm^−1^) of a spinal cord slice (mean thickness = 30 *µ*m) on the region where the birefringence has been measured. (*b*) Measured *H*_2_ map from the polarized signal measured on the spinal cord slice (in this scheme, values are not relevant for optical axes oriented along the horizontal and vertical orientations, see text). (*c*) Deduced birefringence Φ map of the deduced birefringence value. (*d*) Measured *H*_2_ map from the signal measured on a glass coverslip.

Transforming this expression back in the sample plane frame (*X, Y*), the field components are written

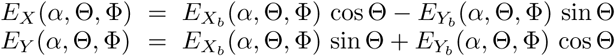

The new intensity *I*_*X*_ (*α*, Θ, Φ) = |*E*_*X*_ (*α*, Θ, Φ)|^2^ dependence can be easily decomposed into circular functions such as:

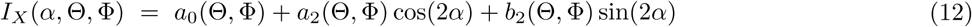

with:

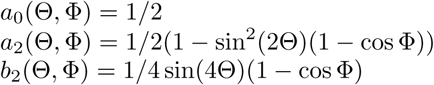

We can define the magnitude and phase of this modulation as:

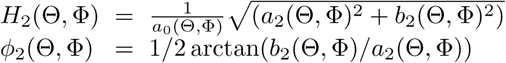

In an isotropic medium, the measured modulation is proportional to cos^2^(*α*), since there is no perturbation of the incident polarization state. The resulting expected values are thus *H*_2_ = 1 and *ϕ*_2_ = 0[*π*/2]. In a birefringent sample 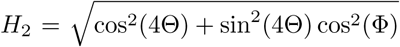 and therefore *H*_2_ is directly related to the birefringence phase shift Φ and the main birefringence axis direction Θ; it is generally lower than 1, except if Φ = 0[*π*], which means no birefringence, or a birefringence value which is an integer number of *π*. The fact that *H*_2_ ≤ 1 is due to the existence of ellipticity, induced by Φ. In order to extract the quantities (Φ, Θ) from the new measured modulation, some operations need to be performed on the modulation parameters (*a*_2_, *b*_2_):

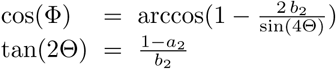

The quantity Φ is thus extracted modulo *π*, while Θ is extracted modulo *π*/2. Note however that due to the measurement geometry (fixed analyzer axis), these quantities cannot be measured for Θ = 0[*π*/2]. A rotation of the analyzer is necessary to disentangle this indetermination.

Values of *H*_2_ calculated from the transmitted signal through the spinal cord sample are depicted in Fig. S3b. *H*_2_ is clearly lower than 1 in many places of the sample, principally concentrated on the myelin sheath contours. The deduced phase shift Φ is shown in Fig. S3c. In this image, the birefringence can be quantified in regions of the myelin sheaths that do not correspond to orientations of the lipids around 0 or 90, for the reason indicated above.

The resulting Φ values range from 20 to 40for the 30 *µ*m (on average) thick slice of the spinal cord sample, which indicates that the sample birefringence is quite large, as also previously reported in the literature. In comparison, the reference glass coverslip *H*_2_ is very close to 1 as can be seen in Fig. S3d.

The found birefringence phase shift extrapolated for a depth of penetration of 30 *µ*m (maximum focus depth used here) is therefore in the range of 20 to 40. The induced polarization distortions have been included in a model of the myelin sheath response (using input (*S*_2_, *S*_4_) values close to the measured ones), following the method described in^39^. The birefringence values are seen to induce a negligible bias of (< 0.05) for *S*_2_, and of 0.05 for *S*_4_. Even though the myelin birefringence is non negligible, the shallow depth penetration used here allows neglecting this effect. However, it means that care has to be taken in case of future investigations at deeper penetration distances.

### C. Molecular order in model lipid membranes

The model membranes, multilamellar vesicles (MLVs) were prepared according to the Bangham et al. method^51^. The phospholipids, dipalmitoylphosphatidylcholine (DPPC) and dioleoylphosphatidylcholine (DOPC) lipids (Avanti Polar Lipids Inc.), were used without further purification. Briefly, thin films of dried lipids were hydrated in pure water (miliQ) above the main phase transition temperature (55C form DPPC or 21C for DOPC), resulting in formation of MLVs varying in size. PR-CARS data were recorded at room temperature (21C), in the spectral region averaged around the CH_2_ symmetric stretching vibration. We considered MLVs having comparable intensity and size in the range from 5 to 15 *µ*m, and imaged both ”filled” and ”non-filled” MLVs that were encountered in the samples (Fig. S4a, b). In order to compare the organization of lipids in both populations, we considered only pixels taken from their border in the equatorial plane. (*S*_2_, *φ*_2_) images exhibit a strong orientational dependence of lipids in the equatorial plane of MLVs as compared to the isotropic aqueous solution, with CH bonds oriented along the membrane contour (Fig. S4c). The highest degree of the lipid order was found for non-filled MLVs made of DPPC lipids, with 〈*S*_2_〉 ≈ 0.40 and 〈*S*_4_〉 ≈ 0.06 (all averages are given over measurements performed over 5 to 10 MLVs). This corresponds to an effective distribution function resembling a Gaussian with an aperture *σ* ≈ 70 (Fig. S4d). Filled MLVs exhibit a higher disorder (Fig. S4d) (〈*S*_2_〉 ≈ 0.29 and 〈*S*_4_〉 ≈ 0.05), most probably because of the lower tension exerted on the outer membranes due to the disorganized inner lipid layers. The measured order can change dramatically with different types of lipids, in particular their degree of saturation. Non-filled MLVs made of DOPC lipids exhibit a lower order with 〈*S*_2_〉 ≈ 0.23, with *S*_4_ almost unchanged (Fig. S4e). This reflects the fact that at room temperature, DPPC is in the gel phase while DOPC is in the liquid disordered phase, with more possible conformational changes that leads to higher disorder^52^.

**FIG. S4.**
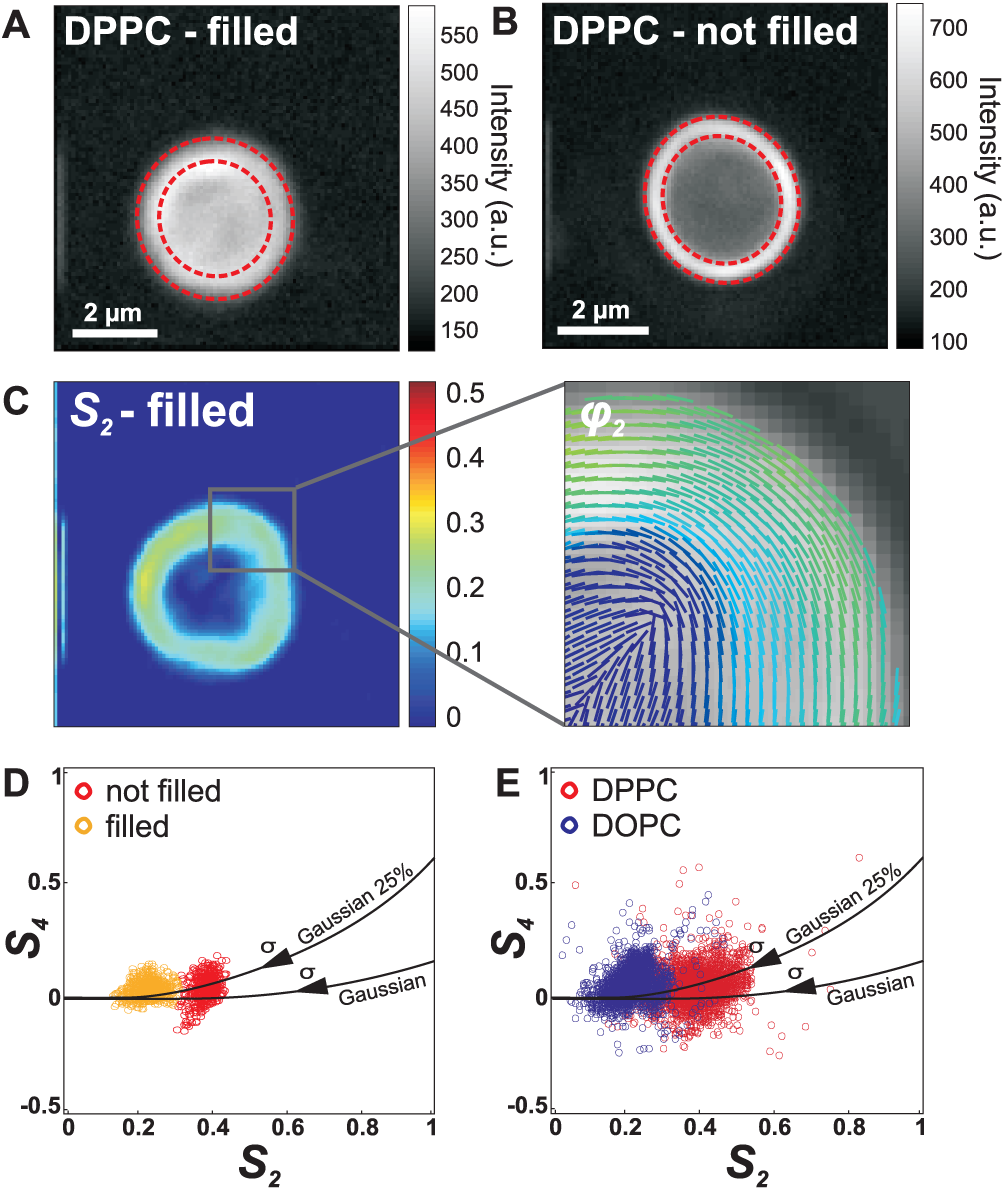
PR-CARS analysis on model multilamellar vesicles (MLVs) (around 2845 cm^−1^). (*a*): intensity CARS image of the ’filled’ MLVs made of pure DPPC. The dashed circles delimitate the region of analysis, emphasizing the outer edge of the MLV. (*b*) similar image for a ’not filled’ DPPC MLV. (*c*) map of *S*_2_ values (left) and corresponding *φ*_2_ image showing the mean orientation of CH molecular bonds (right). (*d*) Comparison of (*S*_2_, *S*_4_) values in ’filled’ and ’not filled’ DPPC MLVs (each point is a pixel taken between dashed circles represented in the intensity images). (*e*) Comparison of (*S*_2_, *S*_4_) values in MLVs made of either pure DPPC or pure DOPC lipids. Continuous lines represent models from a pure Gaussian distribution and a Gaussian distribution superimposed with an isotropic distribution with a proportion of 25%.

